# Solvent Isotope Effect on the Stability of a Heterodimeric Protein

**DOI:** 10.64898/2026.02.12.705670

**Authors:** Rupam Bhattacharjee, Jayant B. Udgaonkar

## Abstract

Protein stability arises from a fine balance between stabilizing forces such as hydrophobic interactions, hydrogen bonding, and ionic interactions, and destabilizing contributions from solvent exposure and electrostatics. Although hydrophobic burial is the dominant driving force for folding, intra-chain hydrogen bonds and ionic interactions modulate stability in context-dependent ways, with effects that vary depending on their location and environment within the protein. Most studies of protein stability have focused on perturbations induced by pH, solvent composition, or mutations in protonated water, leaving the influence of solvent isotopes relatively underexplored. Notably, despite stronger hydrogen bonding in D_2_O, proteins exhibit diverse stability responses upon transfer from H_2_O to D_2_O, suggesting that differential hydration of nonpolar groups plays a key role. Here, the solvent isotope effect on protein stability is investigated using double-chain monellin (dcMN), a β-sheet–rich, two-chain protein with well-characterized folding behavior. By combining conventional equilibrium unfolding measurements with hydrogen-deuterium exchange mass spectrometry (HDX-MS), the stability of wild-type and a less hydrophobic mutant (C42A) dcMN was compared in H_2_O and D_2_O, revealing greater stabilization of the wild-type protein in D_2_O and highlighting the importance of hydrophobic interactions in governing isotope-dependent stability.

## Introduction

The stability of a protein depends on a delicate balance between large stabilizing and destabilizing forces (Dill, 1990; Creighton, 1991). Hydrophobic interactions and backbone hydrogen bonding interactions are the dominant contributors to the stability of the native (N) state of a protein (Makhatadze and Privalov, 1994; Honig and Yang, 1995; Makhatadze and Privalov, 1995; Pace et al., 1996; Pace et al., 2014b). However, the N state is only marginally more stable than the highly heterogeneous unfolded (U) state. In native conditions, the U state is destabilized as the hydrophobic amino acids are exposed to water and cannot form hydrogen bonds with the water molecules, unlike the polar amino acids. The water molecules surrounding the hydrophobic surfaces form water icebergs or clathrates, which are entropically unfavorable (Frank and Evans, 1945; Grdadolnik et al., 2017). Hence, during folding, hydrophobic groups are sequestered away from the water, and become buried inside the interior of the protein, forming tight packing (van der Waals) interactions. On the other hand, the water molecules, after being released from the water icebergs, maximize hydrogen bonding among themselves because there would be a large enthalpic cost otherwise, and the system gains large entropic stabilization upon folding (Kauzmann, 1959). This allows the polypeptide chain to achieve the final three-dimensional folded structure on biologically relevant timescales (Udgaonkar, 2013).

Intra-chain hydrogen bonding also contributes significantly to protein stability (Scholtz et al., 1991; Shirley et al., 1992; Myers and Pace, 1996; Shi et al., 2002; Rose et al., 2006; Bolen and Rose, 2008; Pace et al., 2014a; Pace et al., 2014b), although it is one of the weakest interactions. The energy of a hydrogen bond falls within the range of 3-4 kcal.mol^-1^ (Hagler et al., 1979; Dauber and Hagler, 1980; Fersht, 1999). Proteins contain a large number of hydrogen bonds, and the energies of these hydrogen bonds are additive in nature (Desiraju, 2011), making them significant contributors to protein stability. However, there is debate regarding intra-chain hydrogen bonding facilitating folding as the net number of hydrogen bonds does not change upon folding. Solvent-protein hydrogen bonds break, and intra-chain hydrogen bonds form, raising questions about the effect of hydrogen bonds on protein stability (Klotz and Franzen, 1962; Fersht and Serrano, 1993). Hydrogen bonds can be present either on the surface or in the interior of the protein. As hydrogen bonds are polar in nature, their presence in the interior of the protein may destabilize the protein (Ben-Naim, 1991; Yang et al., 1992; Makhatadze and Privalov, 1993; Privalov and Makhatadze, 1993; Fleming and Rose, 2005; Bolen and Rose, 2008; Rose, 2021). Nevertheless, a stabilizing effect has also been observed (Hebert et al., 1998; Giletto and Pace, 1999; Thurlkill et al., 2006; Gao et al., 2009).

Ionic interactions also play an important role in protein stabilization (Tanford, 1968; Dill, 1990). Salt bridges and unpaired charged side chains are the most common ionic interactions found in proteins. A rational optimization of ionic interactions at the surface of the protein is required for its stability (Loladze et al., 1999; Strop and Mayo, 2000; Perl and Schmid, 2001; Makhatadze et al., 2003; Schweiker et al., 2007; Gribenko et al., 2009). However, ionic interactions at the surface of a protein have also been shown to be destabilizing (Schreiber et al., 1994; Sindelar et al., 1998) or have minimal effect on protein stability (Erwin et al., 1990; Horovitz et al., 1990; Sun et al., 1991). Ionic interactions that are buried in the hydrophobic core of the protein significantly influence the stability of proteins (Matthews, 1995; Giletto and Pace, 1999; Takano et al., 2000; Bosshard et al., 2004). This is because the ionizable residues experience a non-polar environment in the core, and burial in their ionized forms is energetically unfavorable (Rashin and Honig, 1984; Hendsch and Tidor, 1994; Lumb and Kim, 1995; Wimley et al., 1996). As a result, their side chain pK_a_ changes drastically (Westheimer and Schmidt Jr, 1971; Varadarajan et al., 1989; Stites et al., 1991; Loewenthal et al., 1993; García-Moreno et al., 1997; Dwyer et al., 2000; Fitch et al., 2002; Karp et al., 2007; Harms et al., 2009; Aghera et al., 2012). Moreover, this causes the stability of the protein to become pH-dependent (Langsetmo et al., 1991; Khurana et al., 1995; García-Moreno et al., 1997; Dwyer et al., 2000; Harms et al., 2009; Isom et al., 2010; Aghera et al., 2012; Garcia-Seisdedos et al., 2012).

In all of the above studies, stability perturbations of proteins were measured as a function of a change in the pH of the solution, solvent composition, or mutation. These studies utilized protonated water as the solvent. However, the effect of solvent isotopes on stability is not well understood. While the strength of the hydrogen bond increases in D_2_O (O–D…O) compared to H_2_O (O–H…O) (Creswell and Allred, 1962; Benjamin and Benson, 1963; Scheiner and Čuma, 1996), it is not always the case that the stability of protein will be greater in D_2_O than H_2_O. Several proteins, such as RNase A, horse cytochrome *c* and hen egg lysozyme, have been found to exhibit reduced stability in D_2_O (Makhatadze et al., 1995). In contrast, both barstar (Bhuyan and Udgaonkar, 1998) and thioredoxin (Bhutani and Udgaonkar, 2003) show no difference in stability between H_2_O and D_2_O. On the other hand, lactate dehydrogenase (Henderson et al., 1970), staphylococcus nuclease (Antonino et al., 1991), domain 1 of rat CD2 (Parker and Clarke, 1997) and NTL9 (Kuhlman and Raleigh, 1998) have been shown to be stabilized in D_2_O. The extent of stabilization appears to depend on the relative strength of different hydrogen bonds (amide-carbonyl, amide-water and carbonyl-water) in H_2_O and D_2_O (Shi et al., 2002). Moreover, hydrophobic interactions are also influenced in D_2_O as the strength of the bond between water molecules increases (Kresheck et al., 1965). A study on model compounds suggested that the enthalpy of transfer from H_2_O to D_2_O has a significant dependence on the non-polar water-accessible surface area. The enthalpies of transfer of polar groups were found to be zero (Makhatadze et al., 1995). Hence, the differential hydration of buried non-polar groups is the most important factor governing the stability of a protein in H_2_O and D_2_O.

The stability of a protein in isotopic solvent also depends on the nature of its peptide backbone. A protein with either a protonated or a deuterated backbone can exhibit different stabilities in D_2_O. A Study on deuterated RNase A and hen egg lysozyme revealed that they were less stable in D_2_O compared to their protonated forms in H_2_O (Makhatadze et al., 1995). However, when the protonated proteins were dissolved in both D_2_O and H_2_O, they exhibited higher stability in D_2_O than in H_2_O (Huyghues-Despointes et al., 1999; Efimova et al., 2007). Furthermore, the difference in stability in isotopic solvent is independent of the measurement method. Thermal denaturation and chemical denaturation were employed to measure the stabilities of RNase T1 and A in both H_2_O and D_2_O, revealing that both proteins were more stable in D_2_O than in H_2_O (Huyghues-Despointes et al., 1999).

Double chain monellin (dcMN) is an intensely sweet two-chain protein found in the berry of *Dioscoreophyllum cumminsii*. It is a β-sheet rich protein in which a sole α-helix is packed against the five-stranded β-sheet (Figure 1). It is composed of two chains: chain B (β1 – α – β2) is non-covalently attached to chain A (β3 – β4 – β5) (Figure 1). The folding and unfolding of dcMN have been shown to be multistep processes. Several intermediates have been identified on the folding and unfolding pathways of dcMN (Aghera and Udgaonkar, 2012; Bhattacharjee and Udgaonkar, 2021, 2022). However, the equilibrium unfolding transition of wild-type (wt) dcMN appears to be a two-state process when examined using conventional ensemble-averaging probes such as fluorescence and CD (Aghera et al., 2011).

**Figure 1.**
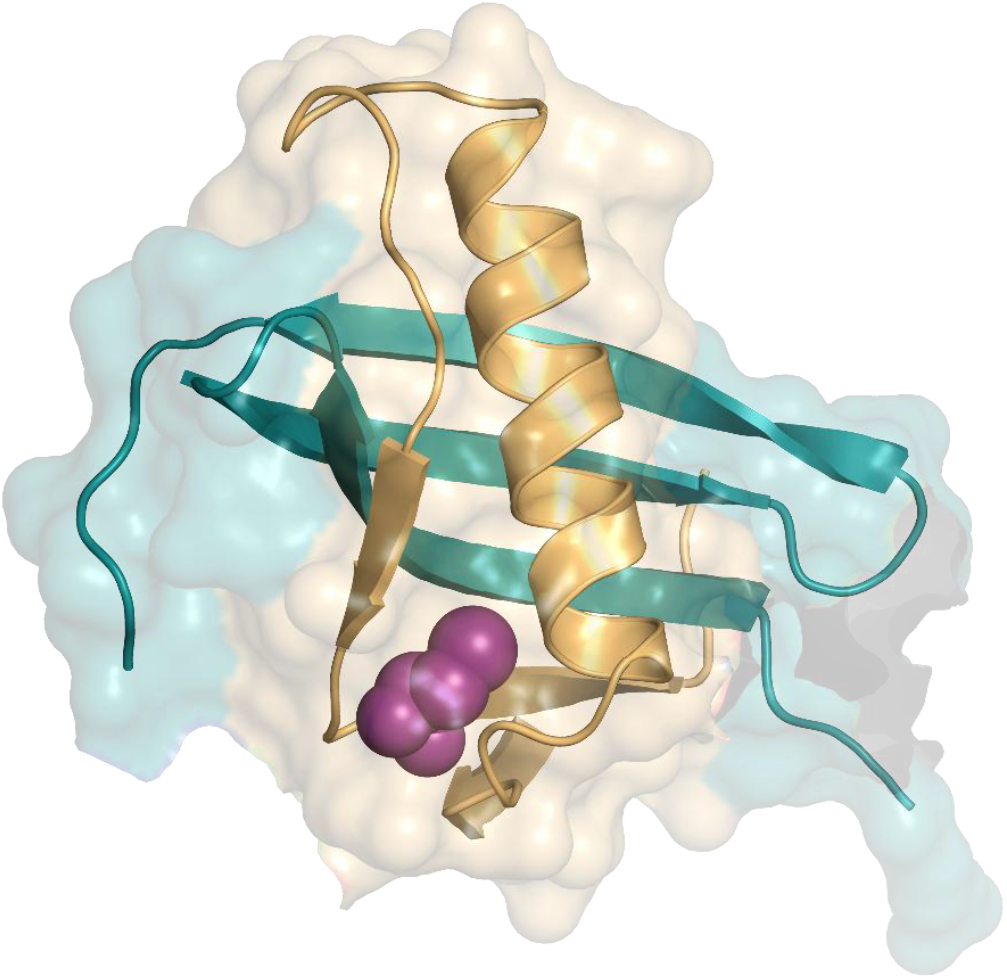
Structure of dcMN. Chains B and A are shown in orange and green, respectively. The cysteine residue at position 42 is shown in purple. The structure was drawn using Pymol and PDB ID 3MON.

In this study, the equilibrium unfolding of protonated wt dcMN is studied along with deuterated wt dcMN to investigate the solvent isotopic effect on the stability of the protein. The equilibrium unfolding transition was also monitored using HDX-MS, which can directly measure the native and unfolded populations, and is likely to identify any small amount of partially protected conformations such as intermediates. A mutant variant of dcMN, C42A dcMN, was also studied under equilibrium conditions due to its reduced hydrophobicity. It was found that the extent of stabilization is higher in the case of wt dcMN compared to C42A dcMN, when the proteins are transferred into D_2_O from H_2_O.

## Materials and Methods

### Protein expression and purification

Double chain monellin was expressed and purified as reported previously (Aghera and Udgaonkar, 2011). The mass and purity of the protein were verified by electrospray ionization mass spectrometry (ESI-MS). The protein was found to be > 95 % pure. Protein concentration was determined by measuring the absorbance at 280 nm, using an extinction coefficient value of 14,600 M^-1^ cm^-1^.

### Buffers and reagents

All the chemicals used in this study were of high purity grade, and were purchased from Sigma. Ultra-pure GdnHCl was obtained from United States Biochemicals, and was of the highest purity grade. All the experiments were carried out at 25 ºC. Sodium phosphate buffer (50 mM) was used at pH 7, and glycine-NaOH buffer (50 mM) was used as the labelling buffer at pH 9, respectively. Ice-cold 100 mM glycine-HCl buffer containing 8 M GdnHCl was used as the quench buffer to stop the labelling reaction. The concentration of GdnHCl in each buffer was determined by measuring the refractive index using an Abbe refractometer. GdnHCl was deuterated by carrying out three cycles of HDX in D_2_O followed by lyophilization. For all D_2_O buffers, the reported pH values were not corrected for any isotope effect.

### Equilibrium unfolding monitored by change in fluorescence, Far-UV CD and Near-UV CD signals

The equilibrium unfolding transition induced by GdnHCl was monitored by measuring fluorescence intensity using the MOS-450 optical system coupled with the SFM-4 mixing module from Biologic. The excitation wavelength was set at 280 nm, and the fluorescence emission was collected at 340 nm using a 10 nm band-pass filter (Asahi Spectra). The final protein concentration was 40 µM. dcMN was incubated in varying concentrations of GdnHCl for 72 h until equilibrium was established before measurement of its optical properties.

GdnHCl- or GdnDCl-induced equilibrium unfolding transitions were also monitored by far-UV CD and near-UV CD measurements on a Jasco J-720 instrument. Far-UV CD measurements were carried out at 222 nm with a 2 mm path length cuvette, while near-UV CD measurements were taken at 270 nm using a 10 mm path length cuvette. For each far-UV CD measurement, the CD signal was averaged over 150 s with a response time of 2 s. For each near-UV CD measurement, the CD signal was averaged over 315 s, using a response time of 2 s.

### Equilibrium unfolding monitored by pulse-labelling HDX

The native protein was deuterated using a pH jump procedure, as described previously (Bhattacharjee and Udgaonkar, 2025). The deuterated native protein was diluted into unfolded buffer (50 mM phosphate buffer prepared in D_2_O, pH 7) to achieve a final GdnDCl concentration of 0-4 M. In all cases, the final protein concentration was 40 µM. The samples were incubated at 25 °C for 72 h until equilibrium was reached. After the equilibrium was attained, a 5 s HX labelling pulse was given by 10-fold dilution into 50 mM Glycine-NaOH buffer (prepared in H_2_O, pH 9) to label the unstructured part of the protein. The HX reaction was quenched by the addition of ice-cold quench buffer, resulting in a final solution pH of 2.8 on ice. The quenched reaction was then incubated on ice for 1 min.

Wild-type dcMN contains a Cys residue at position 42 (in chain B). To prevent inter-chain dimerization through disulfide linkage formation, DTT (prepared in D_2_O) was added to the native protein and buffers, at a final concentration of 2 mM.

### Sample processing for mass spectrometry

After incubation for 1 min on ice, the quenched reaction was desalted using a Sephadex G-25 Hi-trap desalting column from GE, equilibrated with ice-cold distilled water at pH 2.6 (0.1 % formic acid was added), using a Postnova AF4 system. The desalted sample was then injected into the HDX module coupled to a nanoAcquity UPLC (from Waters Corporation), and analyzed using a Synapt G2 HD mass spectrometer (Waters Corporation).

To load the protein on to a C18 reverse phase trap column, water containing 0.05% formic acid was used as the mobile phase at a flow rate of 100 µl/min for 1 min. The two chains of the protein were eluted from the column using a gradient of 35–95% acetonitrile (0.1% formic acid) at a flow rate of 40 µl/min over 3 min. The temperature of the entire chromatography assembly, located inside the HDX module (Waters Corporation), was maintained at 4 ºC to minimize the back exchange during the sample processing (Wales et al., 2008; Walters et al., 2012).

### Data Acquisition by ESI-MS

The source parameters were set to the following values for the ionization of intact protein: capillary voltage, 3 kV; sample cone voltage, 40 V; extraction cone voltage, 4 V; source temperature, 80 ºC and desolvation temperature, 200 ºC.

### Data analysis

#### (a) Analysis of HX data

The protein mass spectrum at each timepoint was generated by combining ∼ 40 scans, each 1 s long, from the elution peak of the total ion count (TIC) chromatogram. Each mass spectrum was then processed further by background subtraction and smoothening using the MassLynx version 4.1 software. From the smoothened spectra, the +7 charge state peak, which exhibited the highest intensity for both chain B and chain A, was selected for further analysis. The signal of the +7 charge state peak was normalized by its total area at each GdnDCl concentration using the Origin software. The mass distributions were fitted to the sum of two Gaussian equations, with each Gaussian distribution corresponding to a specific conformation (U or N). The width(s), height(s) and centroid(s) of the mass distributions were determined. The centroid m/z positions and widths of the mass distributions associated with a particular conformation were allowed to vary within ± 0.1 and ± 0.05, respectively. The fractional area under each Gaussian distribution at a specific GdnDCl concentration represented the fractional population of the corresponding conformation at that GdnDCl concentration. The fractions of unfolded populations at different GdnDCl concentrations were then analyzed using a two-state dimer denaturation model.

#### (b) Fitting of equilibrium unfolding curves to two-state dimer denaturation model

Equilibrium unfolding transitions were fit to a two-state (N ↔ A_U_ + B_U_) dimer denaturation model, in which native protein (N) is in equilibrium with unfolded chain B (B_U_) and chain A (A_U_) (Aghera et al., 2011). The values of the free energy of unfolding in water (ΔG_U_) and the slope of the transition (m value) were obtained from the fit using SigmaPlot 12.

## Results and Discussion

### Both wt and C42A dcMN unfold in an apparent two-state manner in H_2_O and D_2_O at equilibrium

Figure 2 shows fluorescence and CD-monitored equilibrium unfolding transitions for both wt dcMN and C42A dcMN. The equilibrium unfolding curves, monitored by different optical probes, overlap with each other, indicating two-state equilibrium unfolding transitions. Furthermore, these proteins exhibit two-state equilibrium unfolding transitions in both H_2_O and D_2_O solvents. The values for the stabilities, m_U_, and the transition midpoints are listed in Table 1. The stability and midpoint of the equilibrium unfolding curve are found to be higher for C42A dcMN than for wt dcMN in H_2_O (Figure 2, Table 1). One possible reason for this is that the pK_a_ of the thiol group of Cys42 could be higher in the N state compared to in the U state (Aghera et al., 2012). This is because Cys42 is approximately 60% buried in the hydrophobic core of the protein, creating a predominantly hydrophobic microenvironment for the thiol group. As a result, it is destabilized in the core of the protein. Increases in the pK_a_ value of buried polar groups have been observed for many proteins (Khurana et al., 1995; Pinitglang et al., 1997; Dwyer et al., 2000; Tolbert et al., 2005; Harms et al., 2009; Aghera et al., 2012). Additionally, the size of the Cys side chain is larger than that of the Ala side chain, which may hinder the packing of the Cys side chain in the interior of the protein. When Cys42 was mutated to Ala, the polar thiol group was removed, resulting in stabilization of both the core and the protein. However, the m_U_ values were found to be similar (Table 1). This indicates that the change in the solvent-accessible surface area upon unfolding is not significantly different for wt dcMN and C42A dcMN. This is expected because it is highly unlikely that a single amino acid mutation would drastically alter the m-value. For MNEI, when polar residues were mutated to Ala, the stability of the mutant variants was found to be higher than that of wt MNEI (Aghera et al., 2012; Malhotra et al., 2017). The m-values were also found to be unchanged in this case (Aghera et al., 2012; Malhotra et al., 2017).

**Table 1.** Stabilities and m_U_ values of wt dcMN and C42A dcMN in H_2_O and D_2_O.

| Type | | $\Delta G^0$<br>(kcal.mol<sup>-1</sup>) | m-value<br>(kcal.mol<sup>-1</sup> M<sup>-1</sup>) | $C_m$<br>(M) |
| --- | --- | --- | --- | --- |
| Protein | Solvent |  |  |  |
| wt<br>dcMN | H<sub>2</sub>O | 11.8 ± 0.01 | 4.2 ± 0.01 | 1.2 ± 0.01 |
|  | D<sub>2</sub>O | 12.5 ± 0.1 | 4.1 ± 0.02 | 1.5 ± 0.05 |
| C42A<br>dcMN | H<sub>2</sub>O | 12.3 ± 0.05 | 4.1 ± 0.01 | 1.4 ± 0.01 |
|  | D<sub>2</sub>O | 12.5 ± 0.1 | 3.9 ± 0.07 | 1.5 ± 0.05 |

**Figure 2.**
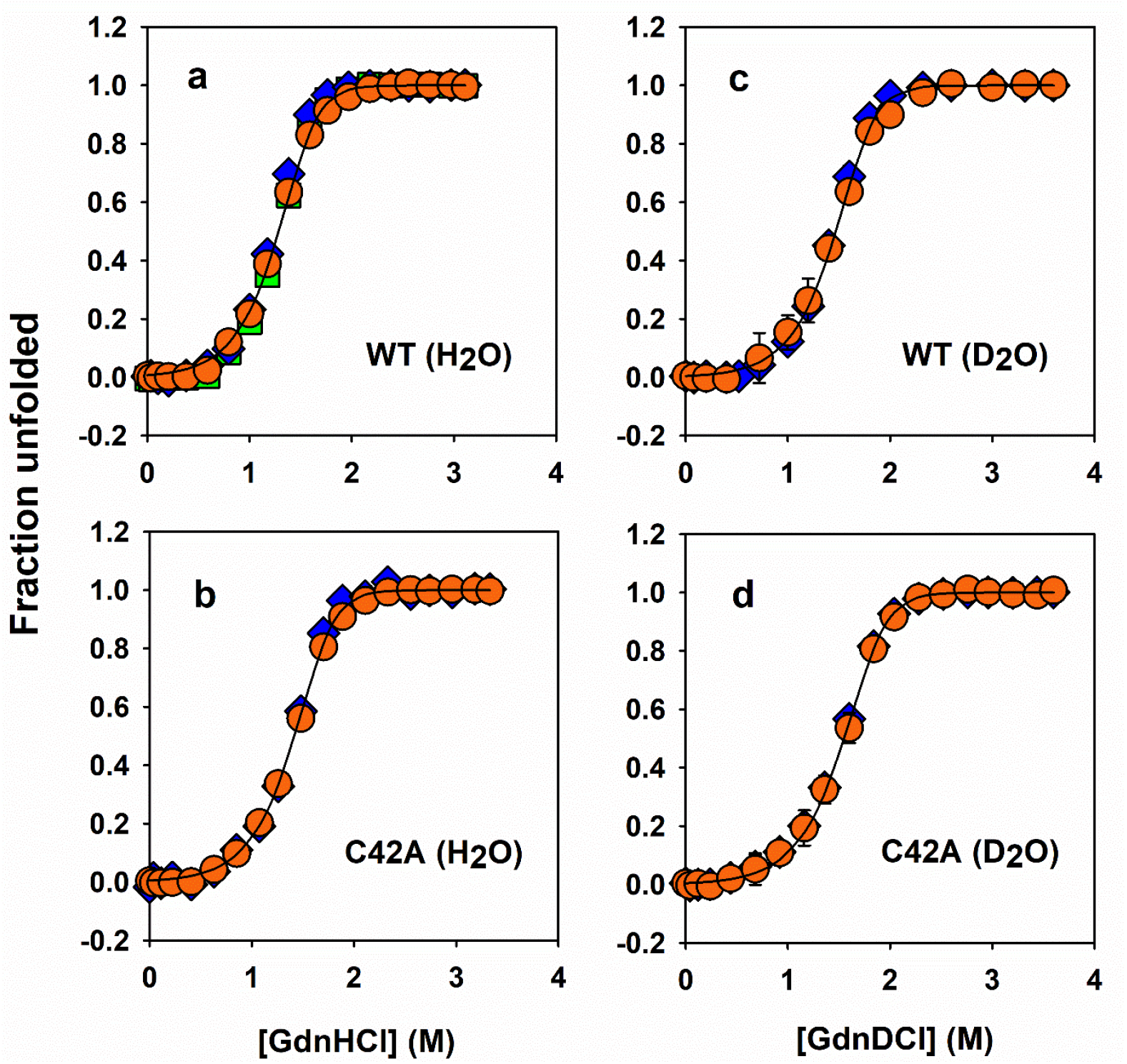
Equilibrium unfolding of wt dcMN and C42A dcMN induced by GdnHCl at pH 7, 25 °C. Panels a and b illustrate the equilibrium unfolding in a protonated solvent for protonated wt dcMN and C42A dcMN, respectively. Panels c and d demonstrate equilibrium unfolding in a deuterated solvent for deuterated wt dcMN and C42A dcMN, respectively. The equilibrium unfolding was determined by monitoring change in intrinsic Trp fluorescence (blue diamonds), far-UV CD (orange circles), and near-UV CD (green squares). The solid lines through the data are nonlinear least-square fits to a two-state (N ↔ A_U_ + B_U_) equilibrium denaturation model. In all experiments, the final protein concentration was 40 μM. The parameters obtained from the fits are summarized in Table 4.1. The error bars represent the spread in the data from two independent experiments.

### Equilibrium unfolding of wt dcMN and C42A dcMN in D_2_O monitored by pulse-labelling HDX-MS

HDX-MS can efficiently identify the populations of different conformations present during a folding/unfolding reaction. To investigate the equilibrium unfolding of deuterated wt dcMN and C42A dcMN, the native and unfolded molecules were pulse-labeled, and their populations were measured. Figures 3a and 3b show the mass distributions of chains B and A of wt dcMN, respectively, at different concentrations of GdnDCl obtained using a 5 s HDX labelling pulse. The mass profiles of both chains fit well to the sum of two Gaussian mass distributions, indicating the presence of two conformations: the more protected N-like conformation, and the less protected U-like conformation (Figures 3c and 3d). The U molecules showed peaks centered at m/z 853.65 and 770.15 for chains B and A, respectively. The N molecules showed peaks centered at m/z 855.95 and 772.75 for chains B and A, respectively, after labelling in the presence of different concentrations of GdnDCl. However, the N molecules retained 1 Da more mass in the absence of the GdnDCl (Figures 3a and 3b) suggesting that the stability of the N conformation decreased in the presence of GdnDCl. A similar observation was made in a previous study, where the N state of dcMN retained fewer deuterium atoms at higher concentrations of GdnHCl (Bhattacharjee and Udgaonkar, 2021).

**Figure 3.**
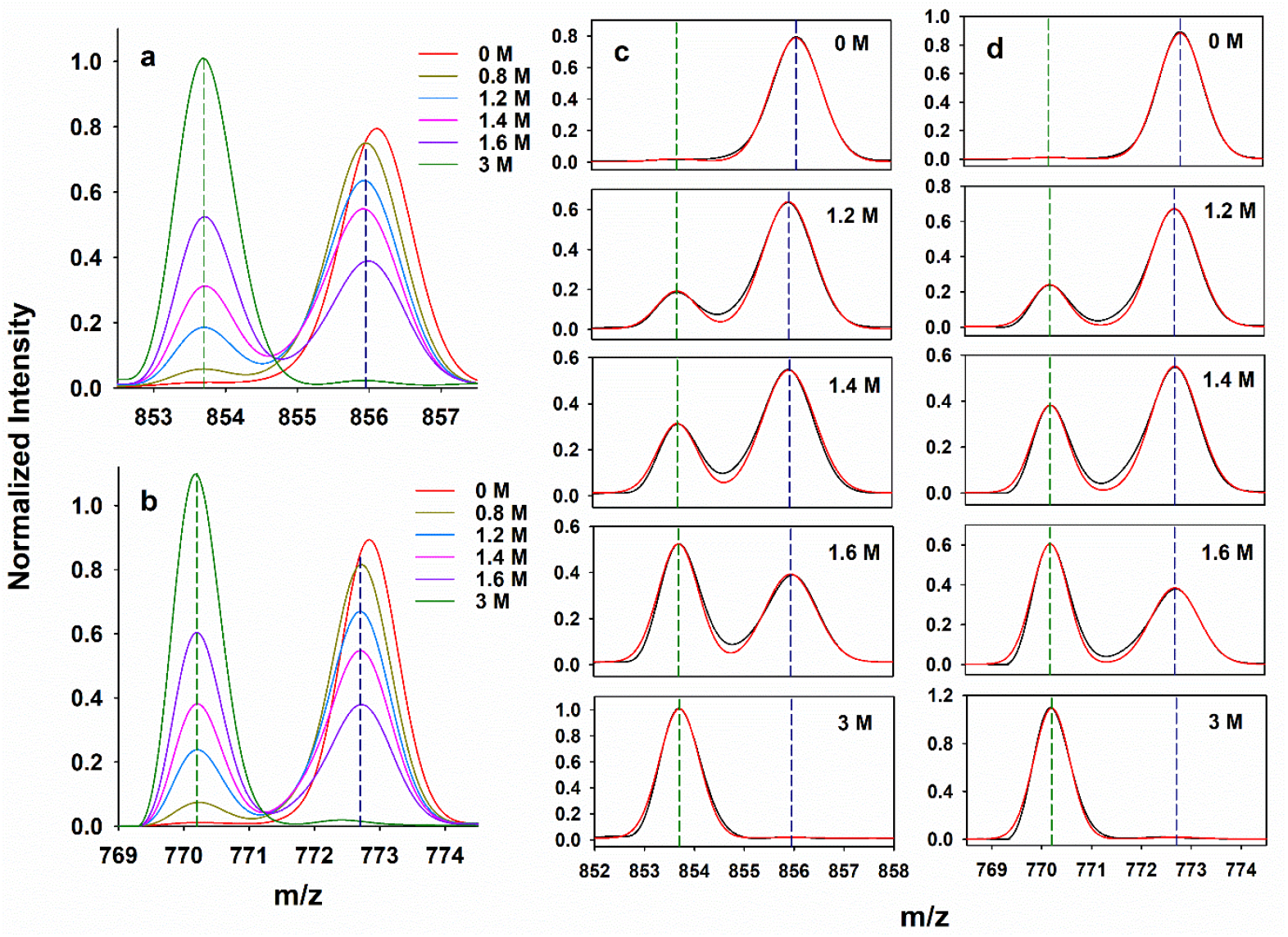
Equilibrium unfolding of deuterated wt dcMN at pH 7, 25 °C monitored using a 5 s HDX labelling pulse at pH 9. Representative mass spectra are shown for chain B (panel a) and chain A (panel b) after labelling in different GdnDCl concentrations. The vertical dashed green and blue lines represent the centres of the mass distributions of the U and N states, respectively. Two-state fits to the mass spectra are shown in panels c and d, for chains B and A, respectively. In each panel, the black line represents the experimentally determined mass profile, and the red line is a fit of the data to the sum of two Gaussian mass distributions. The final protein concentration was 40 μM.

Figures 4a and 4b depict the mass distributions of chains B and A of C42A dcMN, respectively, at different concentrations of GdnDCl, obtained using a 5 s HDX labelling pulse. The mass profiles of both chains were fit to the sum of two Gaussian mass distributions, which indicated the presence of two conformations: the more protected N-like conformation, and the less protected U-like conformation (Figures 4c and 4d). The U molecules showed peaks centered at m/z 849.15 and 770.23 for chains B and A, respectively. The N molecules showed peaks centered at m/z 851.5 and 772.62 for chains B and A, respectively, after labelling in different concentrations of GdnDCl.

**Figure 4.**
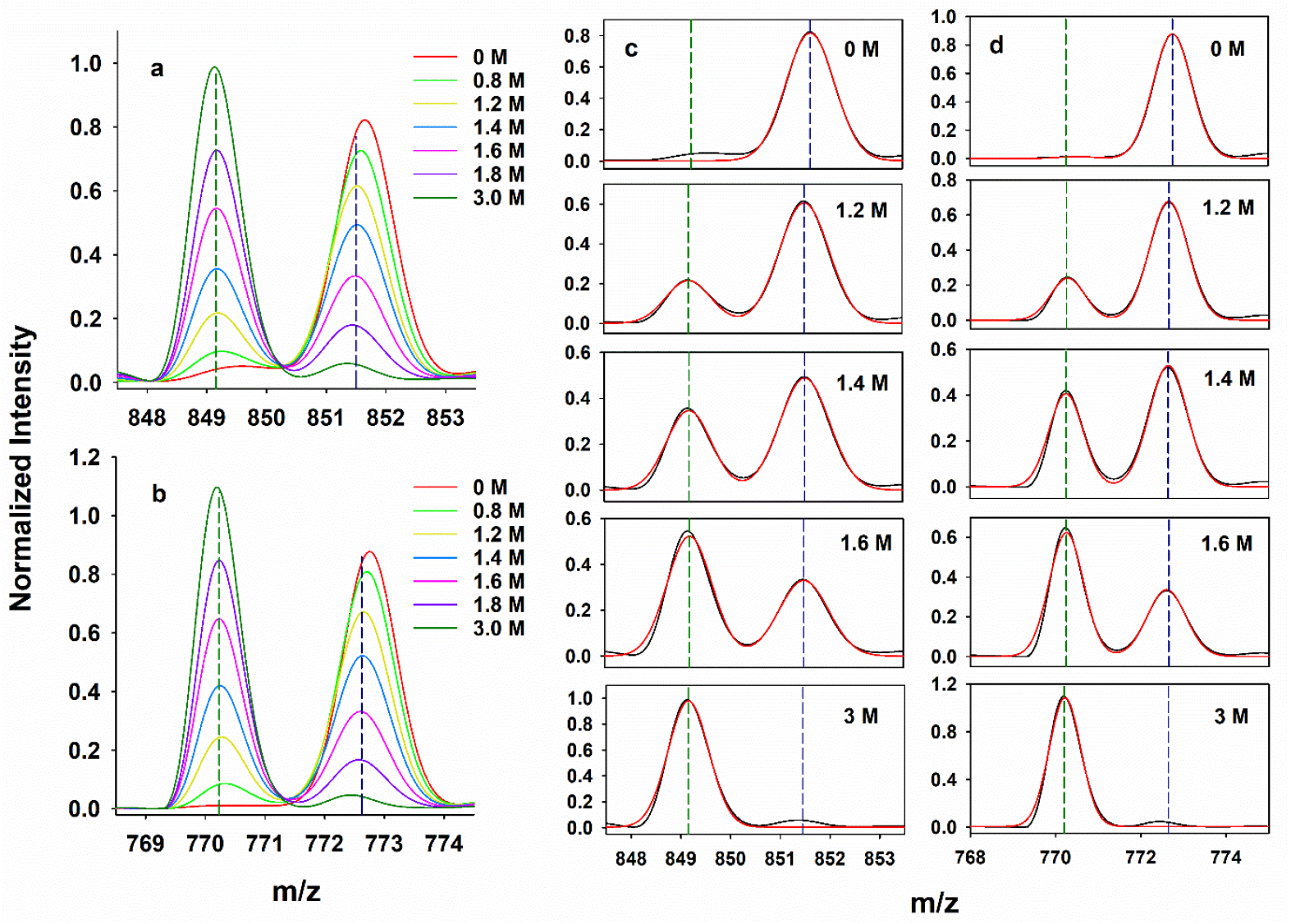
Equilibrium unfolding of deuterated C42A dcMN at pH 7, 25 °C monitored using a 5 s HDX labelling pulse at pH 9. Representative mass spectra are shown for chain B (panel a) and chain A (panel b) after labelling in different GdnDCl concentrations. The vertical dashed green and blue lines represent the centres of the mass distributions of the U and N states, respectively. Two-state fits to the mass spectra are shown in panels c and d, for chains B and A, respectively. In each panel, the black line represents the experimentally determined mass profile, and the red line is a fit of the data to the sum of two Gaussian mass distributions. The final protein concentration was 40 μM.

The mass profiles of chains B and A for both proteins pass through iso-m/z points, suggesting that only two conformations (N and U) are present at any given time, with no intermediates being populated to detectable extents. Thus, the equilibrium unfolding of both proteins appears two-state. The fractional populations of the U-like and N-like conformations at different GdnDCl concentrations were determined from two Gaussian fitting of the HX data. The centroid m/z and width of the mass distributions for the different conformations were kept constant during the fitting. The fractional population of the U conformation was plotted against the concentration of GdnDCl (Figure 5), and was found to overlap with the CD-monitored equilibrium unfolding curve for both wt dcMN (Figure 5a) and C42A dcMN (Figure 5b). This further confirms the two-state behavior of the proteins at equilibrium.

**Figure 5.**
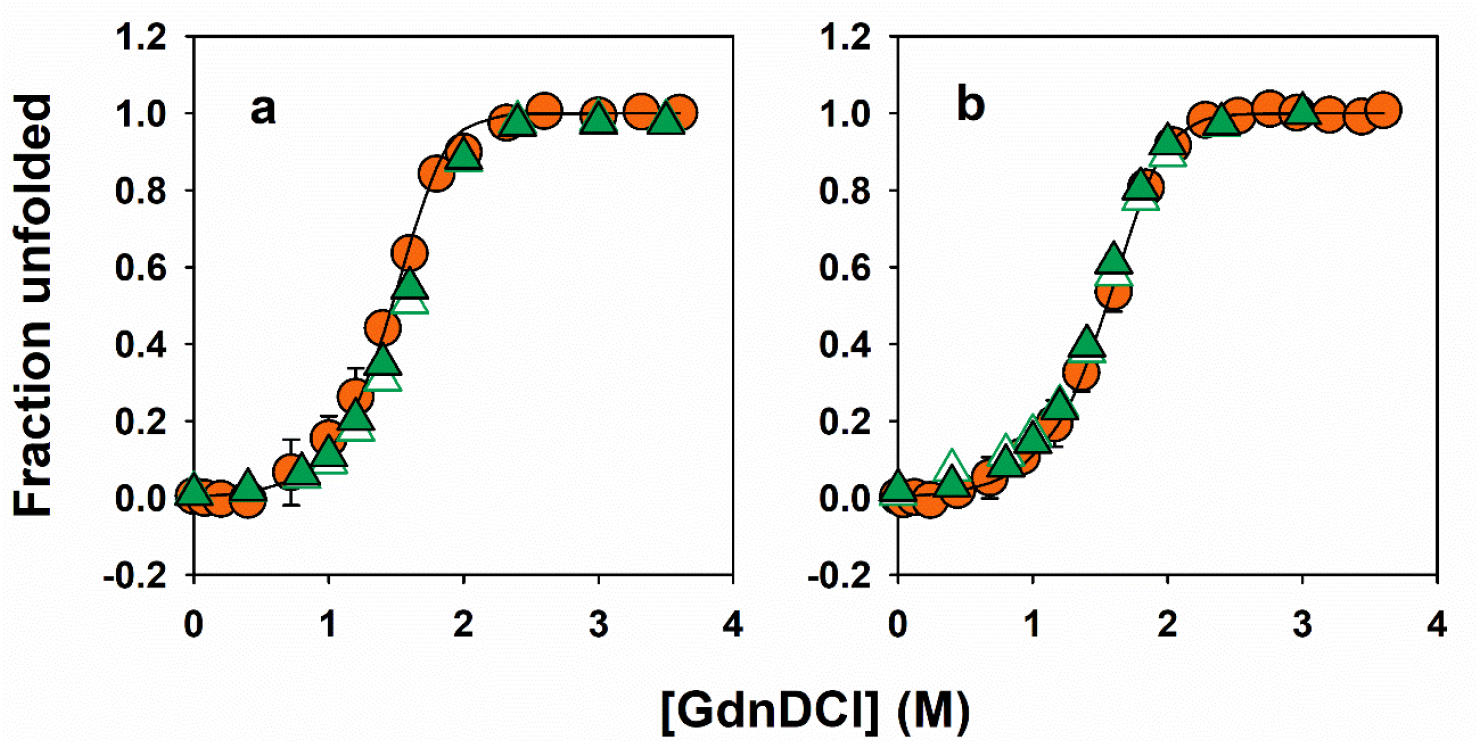
Dependence of the fractional population of U on GdnDCl concentration. Panels a and b show the equilibrium unfolding transitions of wt dcMN and C42A dcMN, respectively, in a deuterated solvent. In both the panels, the circles represent the fractional population of U, which was determined by monitoring the change in ellipticity at 222 nm across different GdnDCl concentrations. The solid lines through the data represent fits of the data to a two-state (N ↔ A_U_ + B_U_) equilibrium denaturation model. The empty and filled triangles correspond to the fraction unfolded obtained from HDX measurements for chains B and A, respectively, which overlaps with the equilibrium unfolding curve obtained by optical measurement. The final protein concentration was 40 μM. The error bars represent the spread in the data from two independent experiments.

### Effect of solvent isotopes on stability

Figure 6a shows the equilibrium unfolding transitions of wt dcMN in both H_2_O and D_2_O. The stability of wt dcMN is 0.7 ± 0.11 kcal.mol^-1^ higher in D_2_O compared to H_2_O (Table 1). This increase is likely associated with the differential hydration of non-polar groups in these two solvents. Previous studies demonstrated that the enthalpies of transfer of polar groups from H_2_O to D_2_O are zero (Makhatadze et al., 1995). This suggests that the hydration energy of polar amino acids is the same in both H_2_O and D_2_O. Therefore, the difference in stability is more likely to originate from the hydration of the non-polar groups, where there would be a higher enthalpic and entropic penalty to hydrate the more hydrophobic surfaces in D_2_O compared to in H_2_O. As a result, the stability of both the N state and the U state would be higher in D_2_O than in H_2_O (Figure 7). However, the extent of destabilization is greater for the U state compared to that of the N state in D_2_O (Figure 7). This is because a greater number of hydrophobic non-polar amino acids are exposed to the solvent in the U state. Hence, the stability of the protein becomes higher in D_2_O than H_2_O (Table 1). The m_U_ value was found to be unaffected upon transferring wt dcMN from H_2_O to D_2_O. Several other studies have also reported higher stabilities of deuterated proteins compared to their protonated forms (Henderson et al., 1970; Antonino et al., 1991; Parker and Clarke, 1997; Kuhlman and Raleigh, 1998). In these studies, the m_U_ values were also found to be not significantly different in D_2_O and H_2_O.

**Figure 6.**
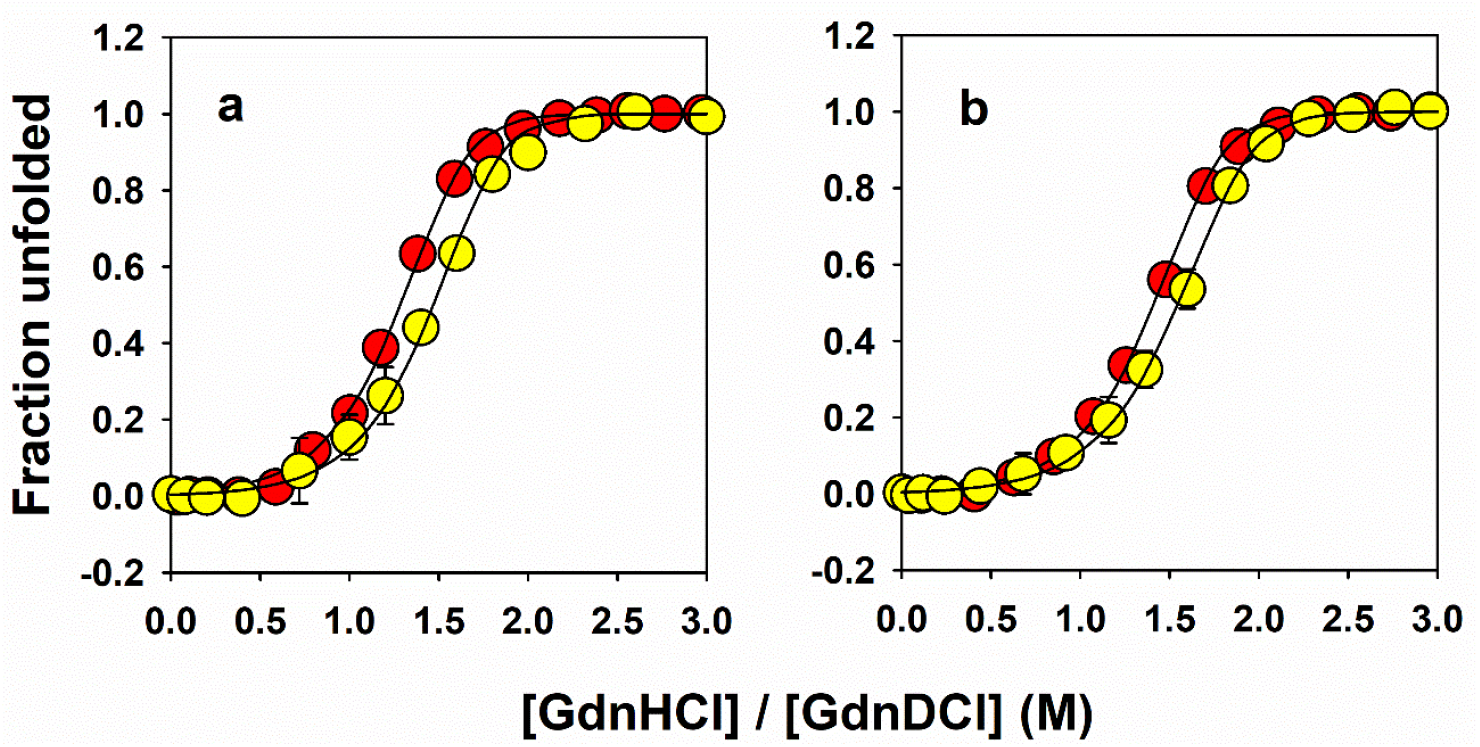
Solvent isotope effect on equilibrium denaturation. Panels a and b show the equilibrium unfolding transitions of wt dcMN and C42A dcMN, respectively. In both panels, the red and yellow circles represent the unfolded fractions for the protonated and deuterated proteins, respectively. Equilibrium unfolding was monitored by measuring the change in ellipticity at 222 nm. The error bars represent the spread in the data from two independent experiments. The final protein concentration was 40 μM. The extent of stabilization resulting from protein deuteration is larger in the case of wt dcMN compared to C42A dcMN (Table 1).

**Figure 7.**
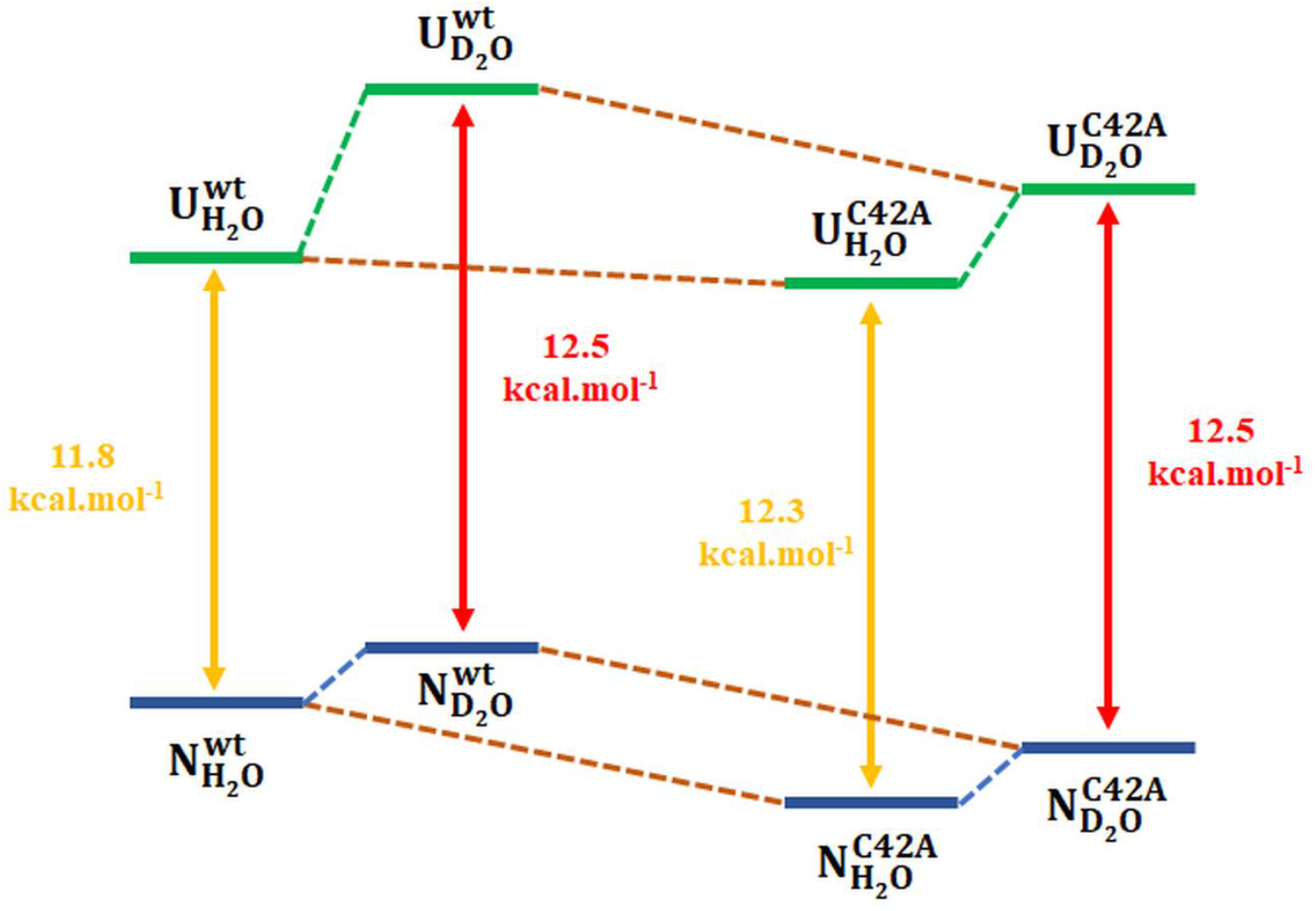
Energy diagram of the N and U states of wt dcMN and C42A dcMN in H_2_O and D_2_O solvents. Both the N and U states are destabilized upon transfer from H_2_O to D_2_O solvents. However, the extent of destabilization is higher in the case of the U states (green dashed line) compared to that of the N states (blue dashed lines). The N state of C42A dcMN is stabilized upon mutation (brown dashed lines). The U state of C42A dcMN is marginally more stable than the U state of wt dcMN in H_2_O (brown dashed line). C42A dcMN is more stable than wt dcMN in H_2_O, but possesses the same stability in D_2_O.

Figure 6b shows the equilibrium unfolding transitions of C42A dcMN in both H_2_O and D_2_O, with an increase in stability of only 0.2 ± 0.15 kcal.mol^-1^ in D_2_O. This result was expected because the hydrophobicity of Ala is slightly lower than that of Cys (Kyte and Doolittle, 1982). Therefore, the hydration of the Cys side chain in the U state is less favorable in D_2_O compared to the Ala side chain. This leads to a destabilization of the U state of wt dcMN to a greater extent than the U state of C42A dcMN in D_2_O (Figure 7). Since the N state of C42A dcMN is more stable than the N state of wt dcMN, the increase in stability in D_2_O is greater for wt dcMN (0.7 kcal.mol^-1^) than for C42A (0.2 kcal.mol^-1^) (Figure 7).

### Protonation state of the protein does not affect the stability in H_2_O and D_2_O

To accurately compare the stability of a protein in H_2_O and D_2_O, it is important to determine the protonation states of buried ionizable residues. Buried ionizable residues may have altered pK_a_ values for their side chains, which can impact the stability of the protein. In the case of D_2_O solvents, the measured D^+^ ion concentration needs to be corrected for the glass electrode solvent isotope artefact using the equation: pD_corr_ = pD_read_ + 0.4, where pD_corr_ and pD_read_ represent the corrected and measured D^+^ ion concentration, respectively (Glasoe and Long, 1960).

For MNEI, the pK_a_ value of the buried Cys42 residue was determined to be 7.5 in the U state, and 9.5 in the N state (Aghera et al., 2012). Since MNEI and dcMN possess identical structures, it can be assumed that Cys42 in dcMN will have a similar pK_a_ value. When the stability of wt dcMN was measured at pH 7 in both H_2_O and D_2_O solvents, Cys42 remained in its protonated form in both solvents. Hence, the observed difference in stability is likely to be due to the solvent isotope effect rather than difference in the protonation state of Cys42.

## Conclusions

In this chapter, the equilibrium unfolding of both wt dcMN and its mutant variant C42A dcMN were investigated in H_2_O and D_2_O solvents. Unfolding was monitored by observing changes in their optical properties and by utilizing HDX. Both proteins exhibited apparently two-state unfolding behavior at equilibrium. Upon transferring from H_2_O to D_2_O solvent, the stabilities of both wt dcMN and C42A dcMN increased. However, the degree of stabilization was found to be higher for wt dcMN compared to C42A dcMN. This difference in stabilization could potentially be attributed to the distinct hydration characteristics of non-polar surfaces in these solvents.

## Acknowledgements

We thank members of our laboratory for discussions. J.B.U is a recipient of a JC Bose National Research Fellowship from the Government of India. This work was funded by the Tata Institute of Fundamental Research and by the Indian Institute of Science Education and Research Pune, as well as by a grant (JBR/2021/000029) from the Science and Engineering Research Board, Government of India.

## Notes

The authors declare no competing financial interest.

